# FieldDino: High-throughput physio-morphological phenotyping of stomatal characteristics for plant breeding research

**DOI:** 10.1101/2024.10.08.617327

**Authors:** Edward Chaplin, Guy Coleman, Andrew Merchant, William Salter

**Affiliations:** School of Life and Environmental Sciences, Sydney Institute of Agriculture, The University of Sydney, Sydney, NSW, Australia; Department of Plant and Environmental Sciences, Faculty of Science, University of Copenhagen, Taastrup, Denmark; The Australian Plant Phenomics Network, The University of Sydney, Narrabri, NSW, Australia

**Keywords:** Stomata, Phenotyping, Wheat, High-throughput, Stomatal Anatomy, Handheld Microscope, Stomatal Conductance, Machine Learning, Plant Breeding, Abiotic stress

## Abstract

Stomatal anatomy and physiology define CO_2_ availability for photosynthesis and regulate plant water use. Despite being key drivers of yield and dynamic responsiveness to abiotic stresses, conventional measurement techniques of stomatal traits are laborious and slow, limiting adoption in plant breeding. Advances in instrumentation and data analyses present an opportunity to screen stomatal traits at scales relevant to plant breeding. We present a high-throughput field-based phenotyping approach, FieldDino, for screening of stomatal physiology and anatomy. The method allows coupled measurements to be collected in <15 s and consists of: (1) stomatal conductance measurements using a handheld porometer; (2) *in situ* collection of epidermal images with a digital microscope, 3D-printed leaf clip and Python-based app; and (3) automated deep learning analysis of stomatal features. The YOLOv8-M model trained on images collected in the field achieved strong performance metrics with an mAP@0.5 of 97.1% for stomatal detection. Validation in large field trials of 200 wheat genotypes with two irrigation treatments captured wide diversity in stomatal traits. FieldDino enables stomatal data collection and analysis at unprecedented scales in the field. This will advance research on stomatal biology and accelerate the incorporation of stomatal traits into plant breeding programs for resilience to abiotic stress.

**Highlight:** Chaplin et al., have developed FieldDino which enables rapid, high-throughput phenotyping of stomatal traits, advancing plant breeding research by integrating streamlined in-field measurements with automated deep learning analysis.

## Background

Stomata control the exchange of gases between the intercellular air spaces of the leaf and the atmosphere. Despite links between stomata, photosynthetic potential and abiotic stress tolerance, stomatal traits have rarely been adopted as targets in plant breeding due to constraints in field measurement at appropriate scales. Here, we outline a coupled high-throughput approach to phenotyping stomatal conductance and stomatal anatomy in the field.

Significant gains in grain yield were made in the second half of the 20^th^ century, largely associated with increased harvest index (grain yield as a proportion of plant biomass) (Porker *et al*., 2020). However, yield improvement has dwindled more recently as realised gains near the theoretical limit in elite populations (Furbank *et al*., 2020). Consequently, the focus of breeders has shifted to radiation use efficiency via increased photosynthetic capacity and efficiency (Parry *et al*., 2011). Stomata play a key role in achieving a higher yield potential (Roche, 2015) and are critical to this increased capacity and efficiency in CO_2_ and H_2_O exchange via dynamic responses to changing environmental conditions. Significant genotypic variation in stomatal conductance has already been identified amongst wheat germplasm (Faralli *et al*., 2024) with high genetic heritability (Rebetzke *et al*., 2013). Development of robust, high-throughput screening approaches could allow us to translate these desirable traits into targets for plant breeding programs.

Previous studies have focussed on characterising either stomatal physiology or stomatal anatomy in isolation. Combining these measures offers considerable advantages for both trait development and modelling. The anatomy of the stomatal cells (including stomatal number and size), is largely fixed after leaf expansion (Mano *et al*., 2023) and is important in determining total pore area (stomatal area index, SAI) and maximum stomatal conductance (*g*_smax_) (Franks *et al*., 2009). Under optimal conditions, lower *g*_smax_ could limit potential CO_2_ assimilation (Wullschleger *et al*., 2002; Harrison *et al*., 2020), whilst under stressful conditions (e.g, drought or heat), a greater *g*_smax_ could lead to excessive water loss (Doheny-Adams *et al*., 2012). In addition, stomatal opening and closure is responsive and regulates instantaneous CO_2_ uptake and water loss relative to changes in the surrounding environment (e.g. light, temperatures, humidity) (Kollist *et al*., 2014; Merilo *et al*., 2014; Nunes *et al*., 2022). Typically, the operating stomatal conductance (*g_sop_*) is significantly lower than the *g*_smax_ (Fanourakis *et al*., 2015; McElwain *et al*., 2016; Ouyang *et al*., 2017). However, *g*_s*op*_ and *g*_smax_ are positively correlated (Franks *et al*., 2009; McElwain *et al*., 2016) and so when considered together, *g*_sop_ and *g*_smax_ (and other environmental data) provide powerful insights into operational efficiencies of leaf gas exchange (Bertolino *et al*., 2019; Busch *et al*., 2024).

Until recently, measurement of stomatal conductance (*g_s_*) has been either (1) fast but lacks precision with porometry or (2) precise but slow with gas exchange. Porometers commonly measure *g_s_* with a humidity or pressure sensor (Bell and Squire, 1981; Monteith *et al*., 1988). They are fast and field portable, however, the accuracy and precision of porometers can vary with several factors, including plant species, water status, environmental conditions and instrument (McDermitt, 1990; Turner, 1991; Fanourakis *et al*., 2016). By contrast, infra-red gas analysers (IRGA) provide more accurate measurements of *g*_s_ (Toro *et al*., 2019), substantially contributing to our understanding of stomatal physiology (Fan *et al*., 2021; Guizani *et al*., 2023; Li *et al*., 2023; Wang *et al*., 2024c), becoming a feature of many laboratories (Salter *et al*., 2018). IRGA’s however are not well suited for high-throughput phenotyping studies, limited to just 10-20 *g*_s_ measurements per hour (Taylor and Long, 2017) and have significant capital cost. Importantly, the development and release of new (circa 2020) handheld leaf porometers, such as the LI-COR LI-600 porometer/fluorometer and the Walz Mini-PAM-II/Porometer, has significantly improved the accuracy of *g*_s_ measurements. With greater reliability under fluctuating environmental conditions, it is now feasible to assess the stomatal conductance of large trials. Deployment in wheat (McAusland *et al*., 2023) and other species (Bheemanahalli *et al*., 2022; Fenstemaker *et al*., 2022; Maestro-Gaitán *et al*., 2022; Caine *et al*., 2023) has rapidly established large datasets.

Image capture of stomatal anatomy has conventionally been similarly low-throughput, time-consuming and costly limiting our ability to screen stomatal traits of large populations. For example, the nail polish ‘peel’ technique is commonly used (Kapadiya *et al*., 2017; Wall *et al*., 2022, 2023), however, suffers from slow cure times, low anatomical precision and contamination with artifacts (Millstead *et al*., 2020) including ‘bubbling’. Recently, a number of published studies have used handheld digital microscopes to capture anatomical images of stomata *in situ* (Sun *et al*., 2021, 2023; Pathoumthong *et al*., 2023) avoiding many of these problems. For example, Pathoumthong et al., (2023) ran a comparative study of the nail polish imprint technique and the use of a handheld digital microscope in wheat, rice and tomato with the latter being faster and less prone to image contamination with air bubbles. Optimisation of this approach including specialised microscope mounts and custom image capture software can allow more extensive anatomical data collection under field conditions.

Post capture, epidermal images must be analysed to extract phenotypic information, such as stomatal number/density, size and/or shape. For plant breeding, thousands of genotypes must be phenotyped – a task that is often time sensitive. Image processing software such as ImageJ (Schindelin *et al*., 2012) has historically been used to aid manual measurement yet this requires significant user input and has little to no automation (Bertolino *et al*., 2019; Caine *et al*., 2019; Dutton *et al*., 2019). Furthermore, such approaches are prone to human error and bias (Casado-García *et al*., 2020). The use of deep learning algorithms and computer vision tools for analysis of microscopy images has developed significantly over the past decade, with deep learning algorithms capable of automatically extracting stomatal features in images of wheat (Gibbs *et al*., 2021; Sun *et al*., 2021, 2023; Pathoumthong *et al*., 2023) and other species (Meeus *et al*., 2020; Ren *et al*., 2021; Zhang *et al*., 2022; Wang *et al*., 2024b,*a*; Yao *et al*., 2024, Preprint). The use of deep learning for stomatal annotation and measurement has significant potential to streamline stomatal anatomy phenotyping, as demonstrated in wheat by Pathoumthong et al. (2023). In the study, they found the data from their YOLOv5 machine learning model was highly correlated with manual measurements for stomata number (R^2^ = 0.99), stomatal size (R^2^ = 0.87) and stomatal aperture (R^2^ = 0.94).

Many models to date have estimated stomatal density with few having the capacity to extract finer morphological details from microscopy images (Gibbs and Burgess, 2024). Furthermore, studies have been limited to only a small handful of genotypes grown under glasshouse conditions with little variation in stomatal anatomy across the dataset. While advancements in computer technology mean that algorithms used for deep learning models will improve quickly, to ensure a high degree of model performance, high quality imagery will always be required. For applicability across a wide range of genotypes, models must be trained using large datasets encompassing diverse plant material imaged in the field. Performance of a more generalised model would also enable invariance to epidermal features, including surface wax and trichomes.

Given recent developments in stomatal phenotyping and associated data analysis, it is now feasible to screen for stomatal physiology and anatomy traits at scales relevant for plant breeding. Our aim was to develop and validate a coupled high-throughput phenotyping approach for stomatal conductance and stomatal anatomy traits. In this study, we describe a three-stage approach: (1) use of a handheld porometer system to collect *in situ* physiological data of *g_s_* and chlorophyll fluorescence, (2) streamlined collection of stomatal anatomy images from the same leaf *in situ* using a handheld microscope with a custom designed 3D printed leaf clip style microscope mount and a custom image capture/microscope control app, and (3) deployment of a deep learning model and computer vision software to automatically segment stomata from the images and analyse their anatomy. We validate this method in large wheat field trials containing 200 diverse genotypes and two irrigation treatments. We describe the approach, present sample output data and outline a future deployment strategy to increase throughput and provide deeper insight into the roles of stomata in the future of crop improvement.

## Methods

### Plant Material

Field plots were established on 24^th^ May 2023 at the University of Sydney I.A. Watson Grains Research Centre in Narrabri, NSW Australia (30.2743°S, 149.8093°E) in two water availability treatments; rainfed and irrigated. 200 genotypes of wheat were sown as part of a broader study focused on drought tolerance of wheat (*Triticum aestivum* L.). The diverse germplasm comprised CIMMYT (International Maize and Wheat Improvement Center) and ICARDA (International Center for Agricultural Research in the Dry Areas) heat tolerant materials imported through the CIMMYT Australia ICARDA Germplasm Evaluation (CAIGE) program. Lines include materials from SATYN (Stress Adapted Trait Yield Nursery), EDPIE (Elite Diversity International Experiment), ESWYT (Elite Selection Wheat Yield Trial), SAWYT (Semi-Arid Wheat Yield Trial), HTWYT (High Temperature Wheat Yield Trials) and the University of Sydney including recombinants from 3 cycles of genomic selection for heat tolerance (including progeny derived from crosses among CIMMYT and ICARDA materials). Three field replicate plots of each genotype arranged in randomised blocks were sown per treatment. Plants were sown in 12 m^2^ plots (2llJ×llJ6 m) at a planting density of 100 plants m^-2^ with five planting rows.

The field site contains predominantly black vertosol cracking clays with high water retention. Minimum tillage was used to maintain soil integrity and moisture. Rainfed plants received 102.4 mm of rainfall and irrigated plants received 197.4 mm of total moisture (rainfall + irrigation) during the growing season. Fertiliser application was consistent across treatments (urea [46% N] 100 kg ha^-1^ and Cotton Sustain [5% N, 10% P, 21% K, 1% Z] 80 kg ha^-1^ pre-planting). The trial was harvested on 26^th^ October 2023.

### Field measurements

For measurements of stomatal physiology and anatomy, one plant was sampled per plot (n = 3 per genotype per treatment). Fully expanded flag leaves were measured at anthesis (Zadok/BBCH stage 48-68) and chosen using a systematic randomised sampling technique, with representative plants of each plot selected from the middle three planting rows, at least 0.5 m from the end of each plot. Data was collected on consecutive days where possible, and all measurements were taken at the same time of day between 09:30 and 15:00 to minimise any diurnal effects on rates of photosynthesis and stomatal conductance. For all leaves sampled, both the upper (adaxial) and lower (abaxial) leaf surfaces were measured.

#### Stomatal conductance

Stomatal conductance and chlorophyll fluorescence of the abaxial and adaxial leaf surfaces was measured using a LI-COR LI-600 porometer/fluorometer (LI-COR Inc., Nebraska, USA) with the following settings: flow rate: 150 µmol s^-1^, flash intensity: 7,000 µmol m^-2^ s^-1^, phase length: 300 ms, ramp amount: 25%, leaf absorptance: 0.8, fraction abs PSII: 0.5, actinic modulation rate: 500 Hz, and integrated modulation intensity: 6.67 µmol m^-2^ s^-1^. The stability was set to the ‘medium’ preset to provide a balance between speed and accuracy. The porometer sensor head was ‘matched’ in the field at the start of the day with an empty chamber and automatically underwent ‘matching’ after every ten measurements throughout the day. Measurements were timestamped and remarks were added to each log.

Abaxial and adaxial measurements were made by placing the sensor head leaf clip on the middle portion of the leaf blade of the main tiller’s flag leaf which was fully sunlit. The leaf was positioned off-centre in the sensor head to avoid clamping the main vein of the leaf. Total stomatal conductance of leaf was obtained by combining the conductance recordings of abaxial and adaxial surfaces.

#### Stomatal anatomy image capture

Following sampling for stomatal conductance, the same leaf was excised from the plant for immediate in-field microscopy imaging of the abaxial and adaxial leaf surfaces using a 200x magnification handheld USB microscope (Dino-Lite 5MP, AM7515MT2A; Dino-Lite, AnMo Electronics Corporation, Taiwan). The 200x magnification was chosen to provide a good balance between sample size for stomatal density and resolution for stomatal size measurements. The microscope was configured with LED brightness = 1, LED axial light = 0 and resolution = 2592 x 1944 pixels.

The reliability of in-field microscope focusing was improved with a custom designed 3D printed leaf clip with a smaller aperture than the cap provided with the microscope. The clip was designed in Tinkercad (Autodesk, Inc., Mill Valley, CA, USA) and files are provided in the FieldDino GitHub repository. Files can be customised for other Dino-Lite microscopes; although STL files for several Dino-Lite models are provided on GitHub. The leaf clip was printed in two parts, the handle/base, and the trigger/microscope mount, which were printed with black and white PETG filament, respectively (Figure 1). Printing the trigger/microscope mount in white also acted as a diffuser for the microscope LED lights. An M5 nut and bolt secure the two parts of the leaf clip together, a small piece of gasket foam is stuck to the surface where the leaf is clamped, and a 38.1 mm extension spring used provide a spring open mechanism after imaging each leaf. Black electrical tape was wrapped around the clear plastic part of the microscope shaft to ensure consistent lighting conditions and prevent external light from hitting the leaf surface. The microscope was connected by USB cable to a laptop computer. Each leaf was imaged by clamping the leaf into the leaf clip, on the same portion of leaf measured with the porometer. Fine focusing on the leaf surface was achieved by adjusting pressure on the trigger with the leaf clamped in the leaf clip. Images of the adaxial and abaxial surface of the leaf were captured before the leaf was discarded. Images were automatically saved to the local hard drive in the order they were captured using DinoXcope software (v2.4.1; Dino-Lite, AnMo Electronics Corporation, Taiwan) in png format.

**Figure 1.**
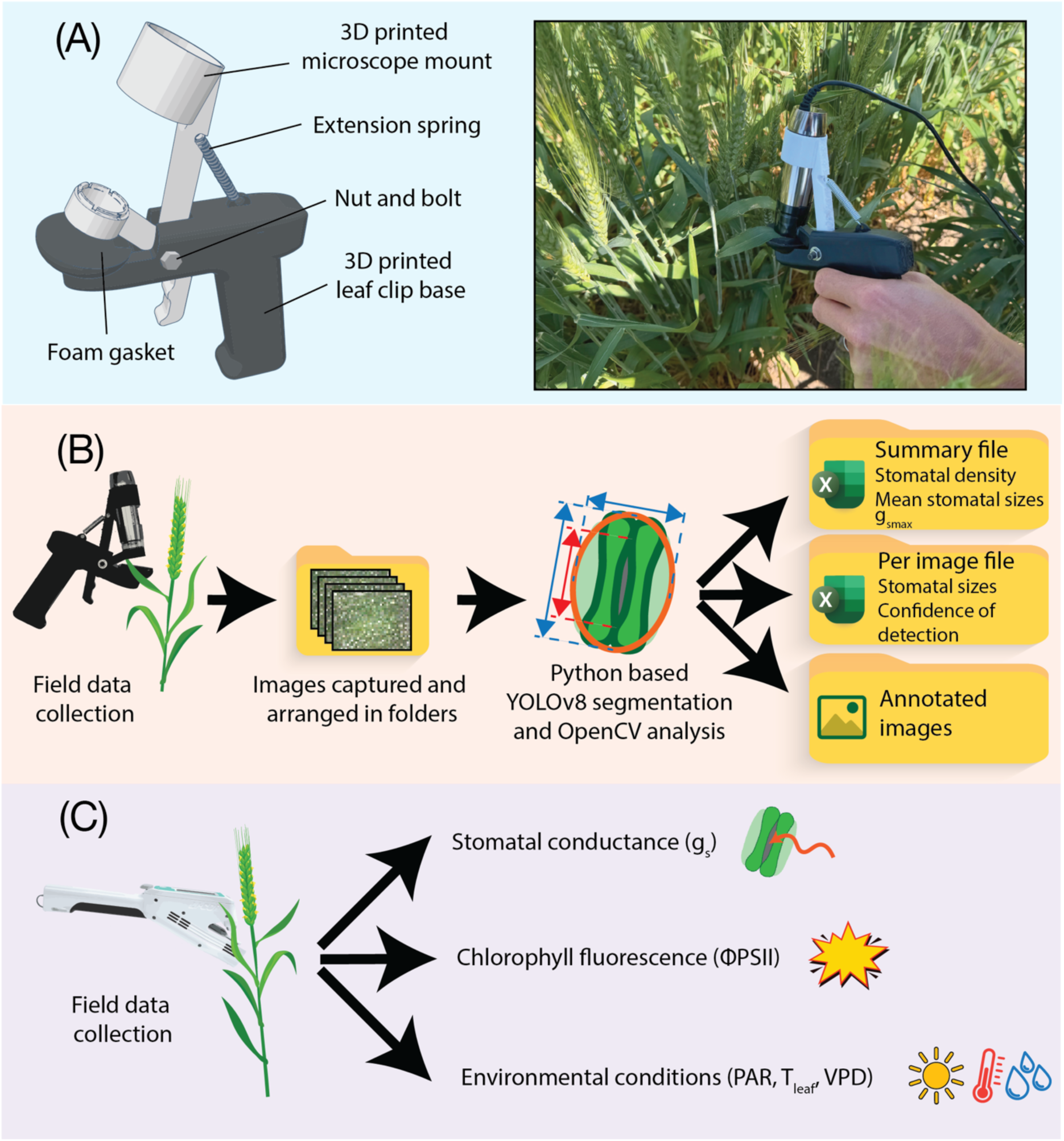
Schematic diagram of the presented stomatal phenotyping method. (A) the 3D printed leaf clip, it’s assembly and in the photo, how the microscope is secured in the leaf clip. (B) and (C) highlight the traits measured by the handheld microscope and porometer, respectively.

#### FieldDino: in-field microscopy app

Following the 2023 field campaign, we developed a custom Python based image capture app optimised for in-field microscopy (Figure 2). The FieldDino app has been tested in subsequent field trials (data not shown) and incorporates improvements over the Dino-Lite software.

**Figure 2.**
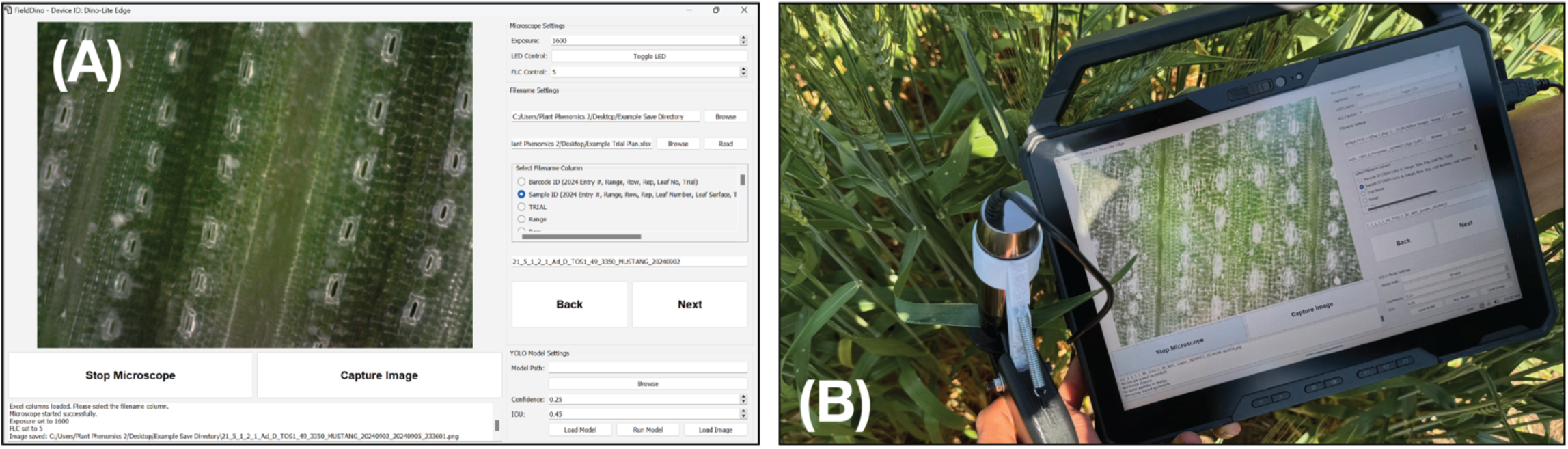
Image capture software. (A) FieldDino custom-designed Python-based app optimised for in-field microscopy. Functionality includes image naming from excel file, a clear video feed with adjustable microscope parameters, easy to use buttons for image capture and ability to run real-time stomatal detection algorithms. (B) handheld digital microscope, custom-made microscope leaf clip and FieldDino app being used in the field for image capture.

The FieldDino app was built with Python and uses a PyQt5 graphical user interface. The software can read image file names from a trial spreadsheet, allowing researchers to efficiently name images according to field/plot locations and a set sampling design. Camera parameters are adjustable and recordable in a metadata text file for consistent record keeping. Additionally, the app features the ability to run real-time deep learning stomata segmentation algorithms using Ultralytics and PyTorch libraries. The app and guidance on its installation and use are available on the FieldDino GitHub repository: https://github.com/williamtsalter/FieldDinoMicroscopy.

### Data analysis

#### Image annotation

A total of 518 full images collected with the Dino-Lite microscope in a single day were used for algorithm development. Images were collected from all 200 genotypes. Stomata were annotated using Roboflow (Roboflow, Inc., Iowa, USA). Polygons for instance segmentation were drawn around the perimeter of the guard cells, pore and subsidiary cells (Figure S1A). Only stomata completely within the image frame were annotated.

To maintain adequate image resolution for the capture the stomatal cell details, whilst enabling effective training, images were tiled into a 4 x 3 grid, producing 12 smaller square sub-images of 640 x 640 from the larger image. Random augmentations were applied to these sub-images prior to export from Roboflow, including random image rotation (± 15°), random brightness (± 15%) and random Gaussian blur of between 0 and 4 pixels with spread measured as the standard deviation in the horizontal and vertical direction from a single pixel. This resulted in a total of 14,928 sub-images in the dataset, split into 13,068 sub-images for training, 1,236 sub-images for model validation and 624 sub-images without augmentation for model testing. The original images (pre-augmentation) are provided as a public dataset on Roboflow: https://universe.roboflow.com/narrabri-plant-physiology-hclvi/fielddino-training-set-200x.

#### Model training and validation

YOLOv8 segmentation models pretrained on the Common Objects in Context dataset were finetuned using the stomata image dataset. All available model sizes were trained, including nano (N), small (S), medium (M), large (L) and extra-large (X). Models were set to train for 1,000 epochs but would cease training if no improvement in performance was made over the previous 100 epochs. The N, S, M, L and X models were trained for 315, 115, 326, 456 and 239 epochs, respectively.

Model performance was compared with precision (P), recall (R), F1 score, mAP@0.5 and mAP@[0.5:0.95] of the mask. Computational efficiency of each algorithm was measured with the Floating Point Operations per Second (FLOPs) and model size, measured in number of parameters.

While accuracy measures how often a model correctly predicts the outcome, precision measures how often the positive class is correctly predicted by the model and thus, how often positive predictions by the model are correct. Recall determines the ability of a model to find all stomata in an image, therefore acting as a measure of whether the model can find all instances of the stomata class. The relationship between the precision and recall metrics is provided by the F1 value (harmonic mean of precision and recall) with the ways in which precision, recall and F1 score are calculated summarised below (Casado-García *et al*., 2020; Padilla *et al*., 2021; Hicks *et al*., 2022; Paniego *et al*., 2022; Pathoumthong *et al*., 2023; Wang *et al*., 2024b).

Precision is defined as:

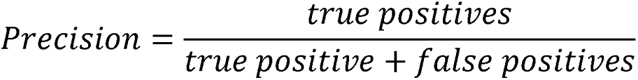

Recall is defined as:

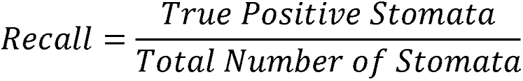

F1 Score is the harmonic mean of precision and recall which is defined as:

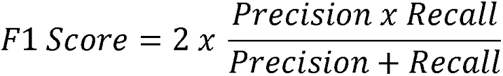

Intersection over Union (IoU) provides an indication of model performance compared with the ground truth annotations, by calculating overlap between the predicted and ground truth regions. Intersection is the area of overlap, representing the region where the prediction has correctly identified the object. Union is the total area which is covered by both the predicted and ground truth segmentation masks (Lin *et al*., 2021; Paniego *et al*., 2022).

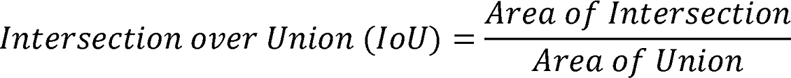

For single-class models average precision (AP) presents a more comprehensive picture of model performance at different IoU thresholds, by computing the area under the precision-recall curve. The commonly used AP@0.5 reflects the AP of the stomata class when used with an IoU threshold of 0.5. A more comprehensive metric, mAP@0.5:0.95 represents the AP over a range of IoU thresholds in increments of 0.05 from 0.5 to 0.95. This therefore assesses the model’s performance at different levels of overlap between the predicted and ground truth boxes (Paniego *et al*., 2022).

Based on these metrics, YOLOv8-M was chosen for stomatal analysis. The model achieved an mAP@0.5 of 97.1% and mAP@[0.5:0.95] of 65.9% (Figure S2).

#### Guard cell – pore length relationship & theoretical anatomical maximal conductance

The anatomical maximum stomatal conductance assuming fully open pores (*g_smax_*) can be calculated from the stomatal dimensions and stomatal density 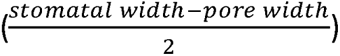 (Franks and Farquhar, 2007; Franks and Beerling, 2009) and the assumption that pore length is approximately half guard cell length (Franks and Farquhar, 2007; Franks and Beerling, 2009). The following equation by Franks and Farquhar (2001) can then be used to calculate *g_smax_*:

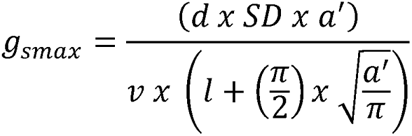

Where: *d* = diffusivity of water in air at 22°C (m^2^ s^-1^), SD = stomatal density (stomata m^-2^), *v* = molar volume of air at 22°C (m^3^ mol^-1^), *l* = pore depth which is equal to the guard cell width at the centre of the stomata based on half the guard cell length (υm), *a^’^*= the mean maximum stomatal pore area which is calculated assuming all stomatal pores are elliptical with the pore length equal to the major axis and the minor axis equal to half the pore length (υm^2^) (Franks and Farquhar, 2001).

To validate the assumption that pore length is approximately half guard cell length as reported by Franks & Beerling (2009) and Franks and Farquhar (2007), prior to running the model, we determined the relationship between the guard cell length and the stomatal pore length. ImageJ was used to measure the length of the guard cell and corresponding pore length of 40 stomata on each of 50 adaxial images and 50 abaxial images where the stomatal pore was clearly visible. This data was then used to plot a regression (R^2^=0.98) to determine the relationship between the two lengths. This enabled us to estimate pore length as 0.543 of the guard cell length (Figure S1B).

#### Measurement of stomata

To determine an appropriate confidence interval with high accuracy and few false-positives, we ran the trained YOLOv8-M segmentation model across all 2,400 images to determine the 5^th^ percentile of confidence intervals. The confidence interval was set to 0.75 and we then ran a custom Python script (available on GitHub), utilising the Ultralytics package for model deployment and OpenCV package for measurements, on all 2,400 images to quantify stomatal traits. The OpenCV script counts the number of stomata per image (stomata mm^-2^), the area of each stomata (µm^2^) and then fits an ellipse around each stomata based on the contours of the polygon produced by the deep learning model (Figure S3). Note, any stomata which were not fully within the image frame were excluded from the analysis to ensure consistency across the dataset. The fitted ellipses were used to estimate guard cell length (µm) and stomatal width (µm). A schematic of the approach and associated metrics is provided in Figure S1A.

#### Analyses and visualisation of data

Data analysis and visualisation were performed with R (R Core Team, 2021) using packages dplyr (Wickham *et al*., 2023), ggplot2 (Wickham, 2016), gridExtra (Baptiste, 2017) and tidyr (Wickham *et al*., 2024). Note that we have not performed any contextual statistics on this data as yet, as this data is part of a larger dataset and this is not the primary focus of our work here. We have provided qualitative descriptions of observable trends in the data.

## 3 Results

To validate the deployability of our coupled phenotyping method at scales relevant to plant breeding, we collected data across a large, replicated field trial on 200 diverse wheat genotypes. In total, we collected 2,400 coupled measurements of stomatal conductance/chlorophyll fluorescence and stomatal anatomy across 5 days, with all measurements collected between 09:30 and 15:00 each day. Each measurement took < 15 s. The total time per measurement increased to 1 min with the inclusion of leaf selection and movement plot to plot; however, these are experiment/trial design specific.

### 3.1 Stomatal conductance and chlorophyll fluorescence

Independent of treatment and genotype, the adaxial stomatal conductance tended to be greater than the abaxial conductance. With respect to water availability, *g*_sop_ tended to be greater under irrigated conditions compared to rainfed. The mean abaxial *g*_sop_ was 0.08 mmol m^-2^ s^-1^ and 0.06 mmol m^-2^ s^-1^ for irrigated and rainfed treatments, respectively, while the mean adaxial *g*_sop_ was 0.20 mmol m^-2^ s^-1^ and 0.14 mmol m^-2^ s^-1^ for irrigated and rainfed treatments, respectively. *g*_sop_ was also variable across the 200 genotypes (Figure 3) with median *g*_sop_ ranging from 0.13 mmol m^-2^ s^-1^ to 0.19 mmol m^-2^ s^-1^ and 0.01 mmol m^-2^ s^-1^ to 0.13 mmol m^-2^ s^-1^ for the abaxial surface under irrigated and rainfed conditions, respectively. The median conductance of the adaxial surface ranged from 0.06 mmol m^-2^ s^-1^ to 0.39 mmol m^-2^ s^-1^ and 0.03 mmol m^-2^ s^-1^ to 0.30 mmol m^-2^ s^-1^ for the adaxial surface under irrigated and rainfed treatments, respectively.

**Figure 3:**
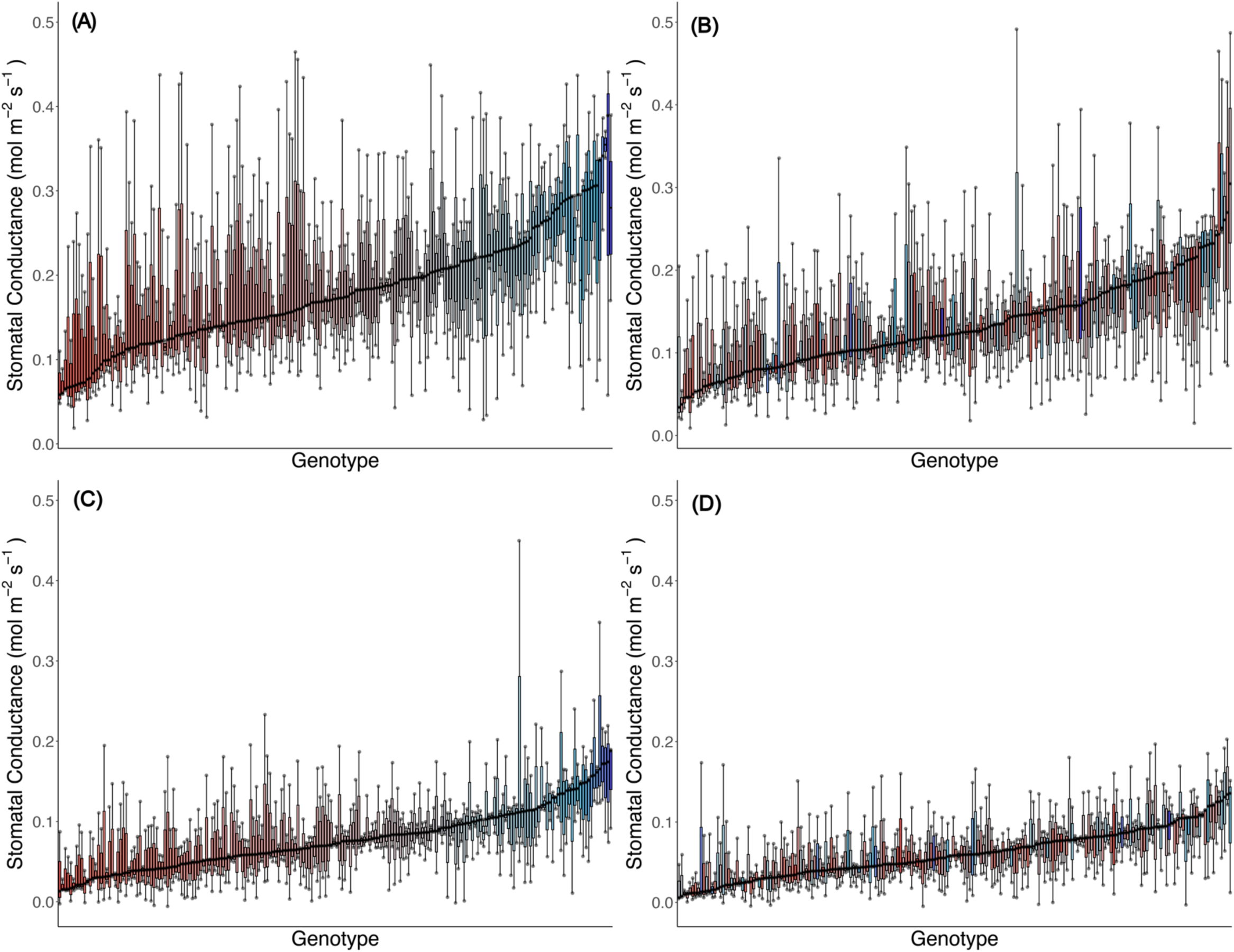
Genotypic distribution of operational stomatal conductance (*g_sop_*) across 200 wheat genotypes under two watering treatments. (A) Adaxial surface irrigated; (B) adaxial surface rainfed; (C) abaxial surface irrigated; and (D) abaxial surface rainfed. Genotypes are ranked by median operational stomatal conductance (*g_sop_*). Colour assigned to each genotype based on irrigated stomatal conductance. Thick horizontal lines within boxes indicate the median and boxes indicate the upper (75%) and lower (25%) quartiles. Whiskers indicate the ranges of the minimum and maximum values. Points indicate individual measurements.

The quantum yield of electron transfer at photosystem II (ΦPSII) tended to be lower when water was more limited (Figure 4) despite the range of values being similar across the treatments. The median abaxial ΦPSII was 0.61 mmol m^-2^ s^-1^ and 0.46 mmol m^-2^ s^-1^ for irrigated and rainfed treatments, respectively, while the median adaxial ΦPSII was 0.58 mmol m^-2^ s^-1^ and 0.46 mmol m^-2^ s^-1^ for irrigated and rainfed treatments, respectively. Genotypes varied in ΦPSII (Figure S4) with median values for the adaxial surface ranging from 0.17 mmol m^-2^ s^-1^ to 0.73 mmol m^-2^ s^-1^ and 0.16 mmol m^-2^ s^-1^ to 0.70 mmol m^-2^ s^-1^ for the rainfed and irrigated treatments, respectively. For the abaxial surface, median ΦPSII values ranged from 0.17 mmol m^-2^ s^-1^ to 0.73 mmol m^-2^ s^-1^ and 0.15 mmol m^-2^ s^-1^ to 0.74 mmol m^-2^ s^-1^ for the two treatments, respectively. Whilst the porometer/fluorometer also reported electron transport rate, we found this to be noisy on the days we recorded measurements, due to patchy fast-moving clouds causing high variability in the light environment. This data was discarded however, we do consider this in our discussion.

**Figure 4:**
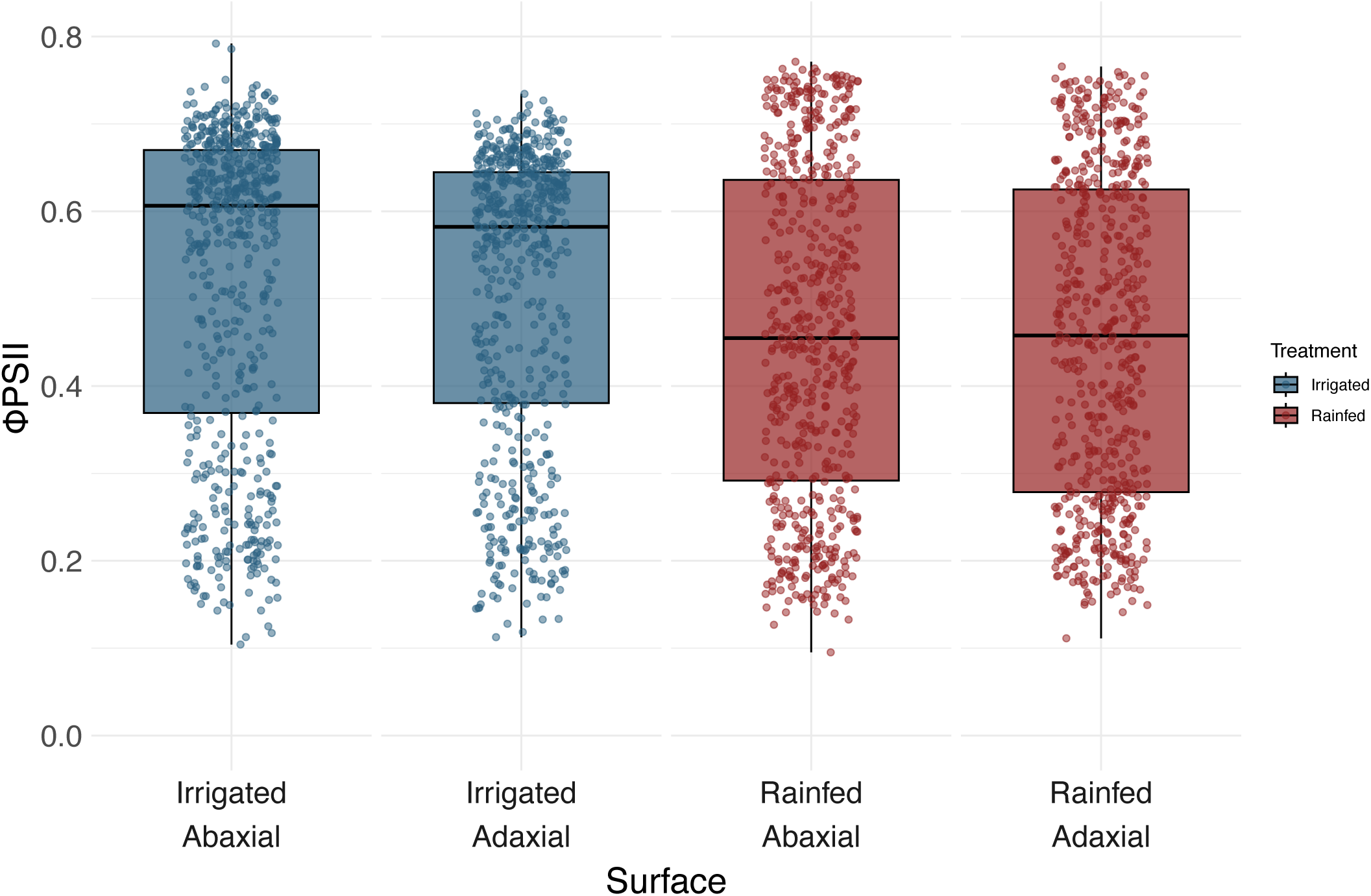
ΦPSII of the abaxial and adaxial leaf surfaces of wheat under irrigated and rainfed conditions. Thick horizontal lines within boxes indicate the median and boxes indicate the upper (75%) and lower (25%) quartiles. Whiskers indicate the ranges of the minimum and maximum values. Points indicate individual measurements.

### 3.2 Stomatal Anatomy

Microscopy images of the leaf surface were collected from the same leaves used for stomatal conductance measurements. Representative images of stomatal anatomy from the abaxial and adaxial leaf surfaces before and after stomatal detection using the deep learning model are shown in Figure 5.

**Figure 5:**
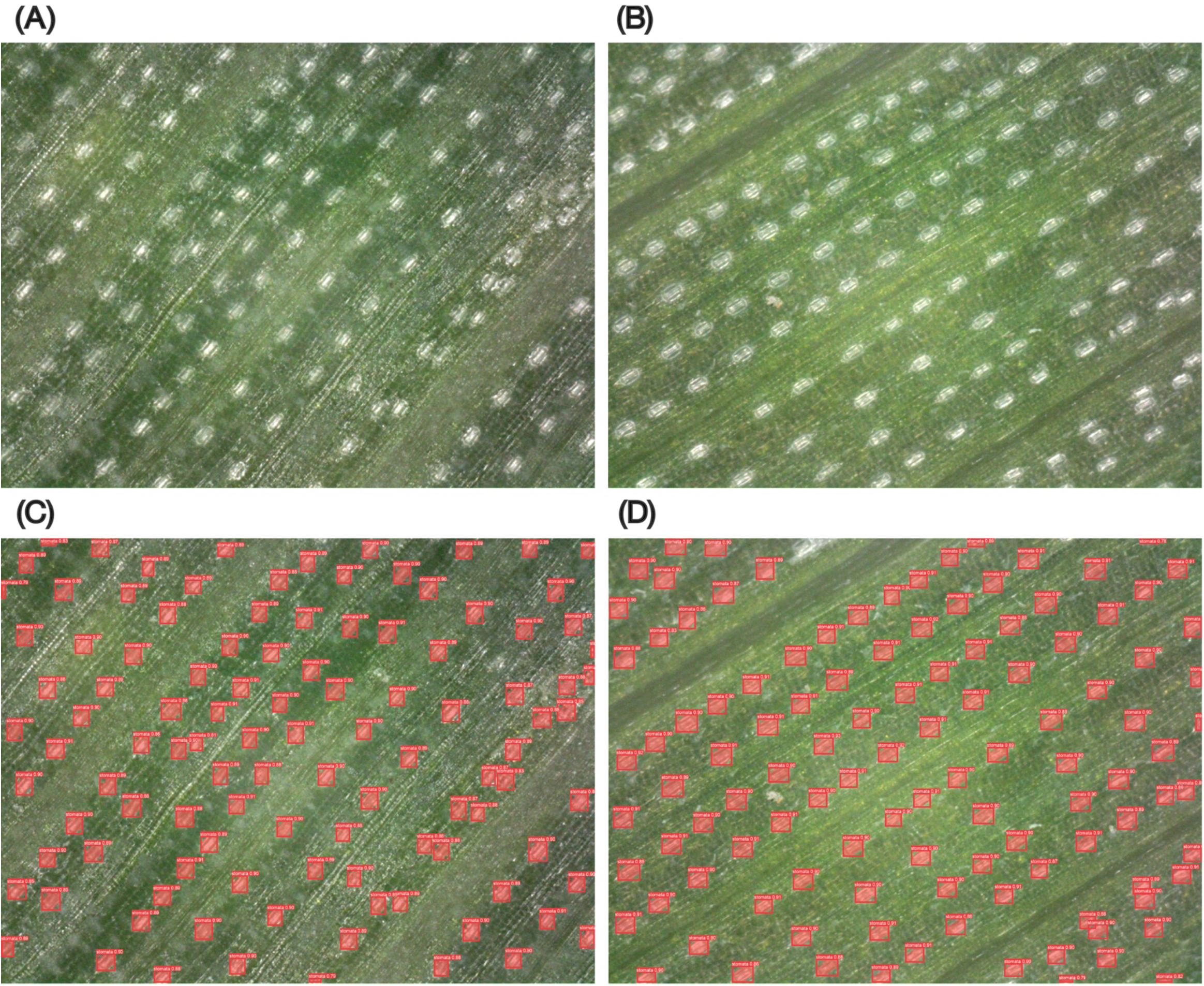
Representative images of stomatal anatomy collected in situ using handheld digital microscope. (A) and (C) abaxial and (B) and (D) adaxial leaf surfaces. Custom leaf clip mount was used with microscope to aid with image focussing. (A) and (B) show raw image collected using microscope, (C) and (D) show image with automatically labelled stomata using the deep learning model we trained.

Under irrigated conditions, guard cell length averaged 68.39 µm and 68.55 µm for the abaxial and adaxial surfaces, respectively. Guard cell length tended to be smaller under rainfed conditions with averages of 64.30 µm and 66.04 µm for the abaxial and adaxial surfaces, respectively (Figure 6A).

**Figure 6:**
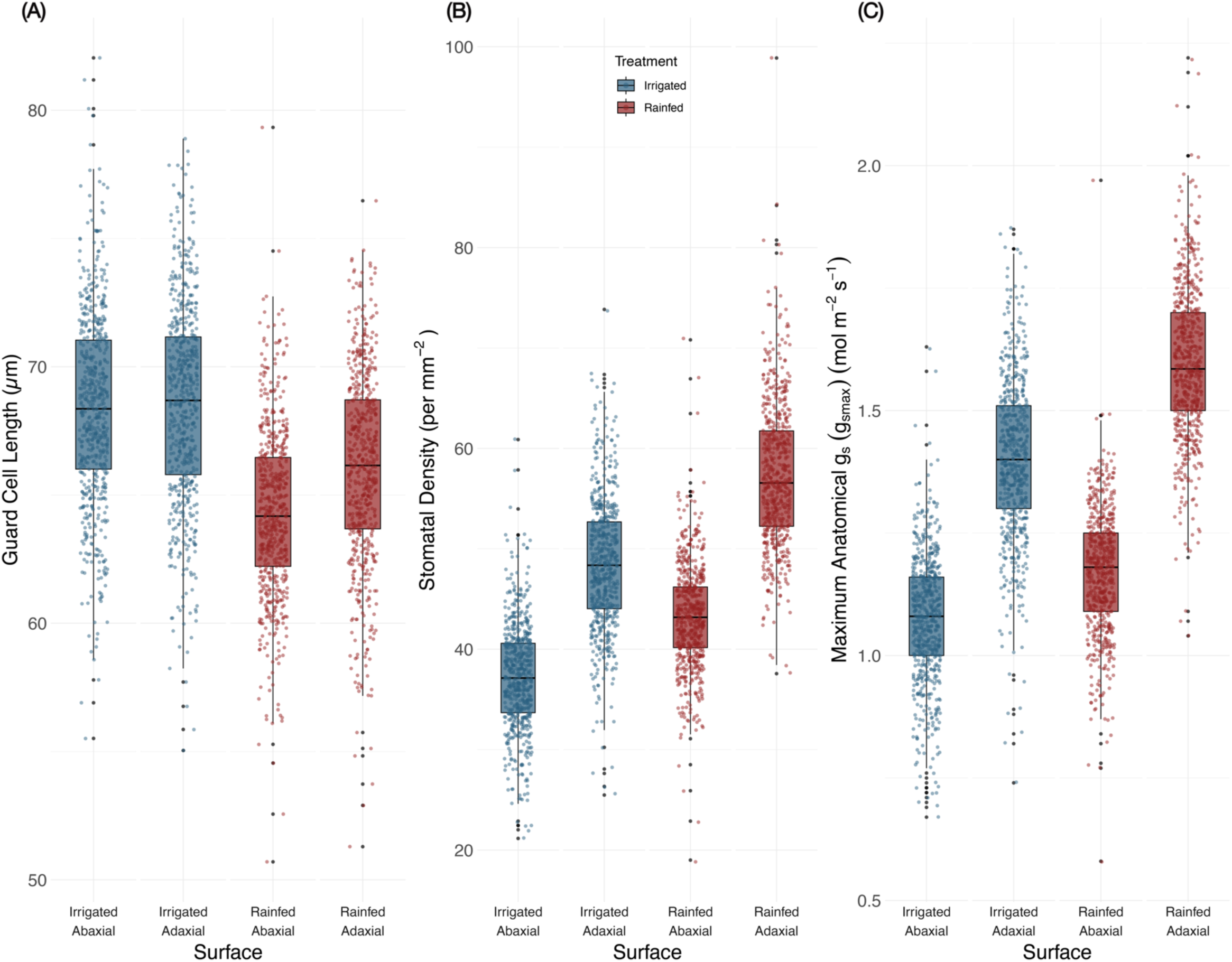
Stomatal anatomy traits of wheat under irrigated and rainfed treatments. (A) Guard cell length, (B) stomatal density and (C) maximal anatomical stomatal conductance, *g_smax_*. Thick horizontal lines within boxes indicate the median and boxes indicate the upper (75%) and lower (25%) quartiles. Whiskers indicate the ranges of the minimum and maximum values. Points indicate individual measurements.

When stomatal density was visualised, we found that the adaxial surface of the leaf generally had a greater stomatal density compared with the abaxial leaf surface. The mean stomatal density was 50.31 per mm^2^ and 42.87 per mm^2^ for the adaxial and abaxial surfaces, respectively. We also noted that the rainfed plants tended to have a higher stomatal density than the irrigated plants (Figure 6B). We observed genotypic variation in stomatal density across the 200 genotypes used for the trial with median stomatal densities for the adaxial surface ranging from 35.8 per mm^2^ to 62.6 per mm^2^ and 42.8 per mm^2^ to 80.7 per mm^2^ for the irrigated and rainfed treatments, respectively. By contrast, for the abaxial surface, the median stomatal density ranged from 25.9 per mm to 49.7 per mm^2^ to 33.7 per mm^2^ to 63.5 per mm^2^ for the two treatments, respectively. We visualised the correlation between stomatal size and density and observed a negative correlation with stomatal size decreasing with increasing stomatal density (Figure 7). We also found genotypic variation across the 200 genotypes used for the trial with regards to both stomatal density and stomatal size.

**Figure 7:**
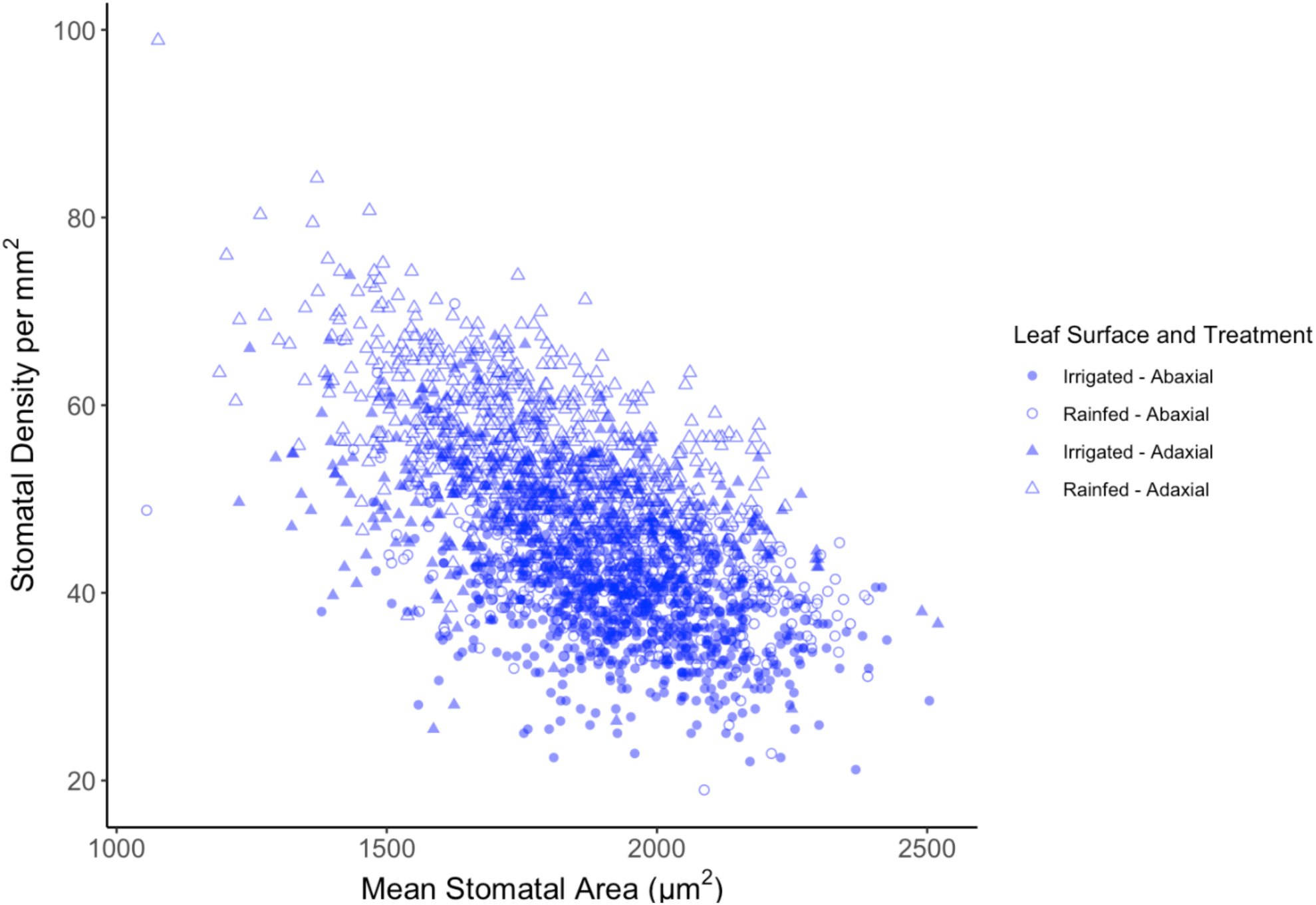
Relationship between stomatal density and stomatal area. Shaded markers represent irrigated plants and unshaded markers represent rainfed plants. Circular markers represent abaxial leaf surface and triangular represent adaxial leaf surface.

*g_smax_* was calculated based on stomatal anatomy, ranging from 0.58 mol m^-2^ s^-1^ to 2.22 mol m^-2^ s^-1^ with the adaxial surface *g_smax_* tending to be larger than abaxial *g_smax_* (Figure 6C). Genotypic variation in *g_smax_* was observed across the 200 genotypes used in the trial with median *g_smax_* values for the adaxial surface ranging from 1.12 mol m^-2^ s^-1^ to 1.73 mol m^-2^ s^-1^ to 1.20 mol m^-2^ s^-1^ to 1.86 mol m^-2^ s^-1^ for the irrigated and rainfed treatments, respectively. For the abaxial surface, median *g_smax_* values ranged from 0.78 mol m^-2^ s^-1^ to 1.29 mol m^-2^ s^-1^ to 0.90 mol m^-2^ s^-1^ to 1.48 mol m^-2^ s^-1^ for the two treatments, respectively.

### 3.3 Combined stomatal physiology and anatomy traits

The coupled physiology-anatomy approach we deployed allowed the stomatal conductance operating efficiency to be calculated. The operating efficiency (*g_s_*_op_/*g_smax_*) tended to be lower under rainfed conditions compared with irrigated conditions and the median operating efficiency of the adaxial surface was higher than the abaxial surface. Variation in the *g_s_* operating efficiency was observed across the 200 genotypes (Figure 8) with median adaxial surface operating efficiencies of 0.03 to 0.25 and 0.01 to 0.17 for the irrigated and rainfed treatments, respectively. For the abaxial surface, the median operating efficiency ranged from 0.02 to 0.29 and 0.01 to 0.17 for the two treatments, respectively.

**Figure 8:**
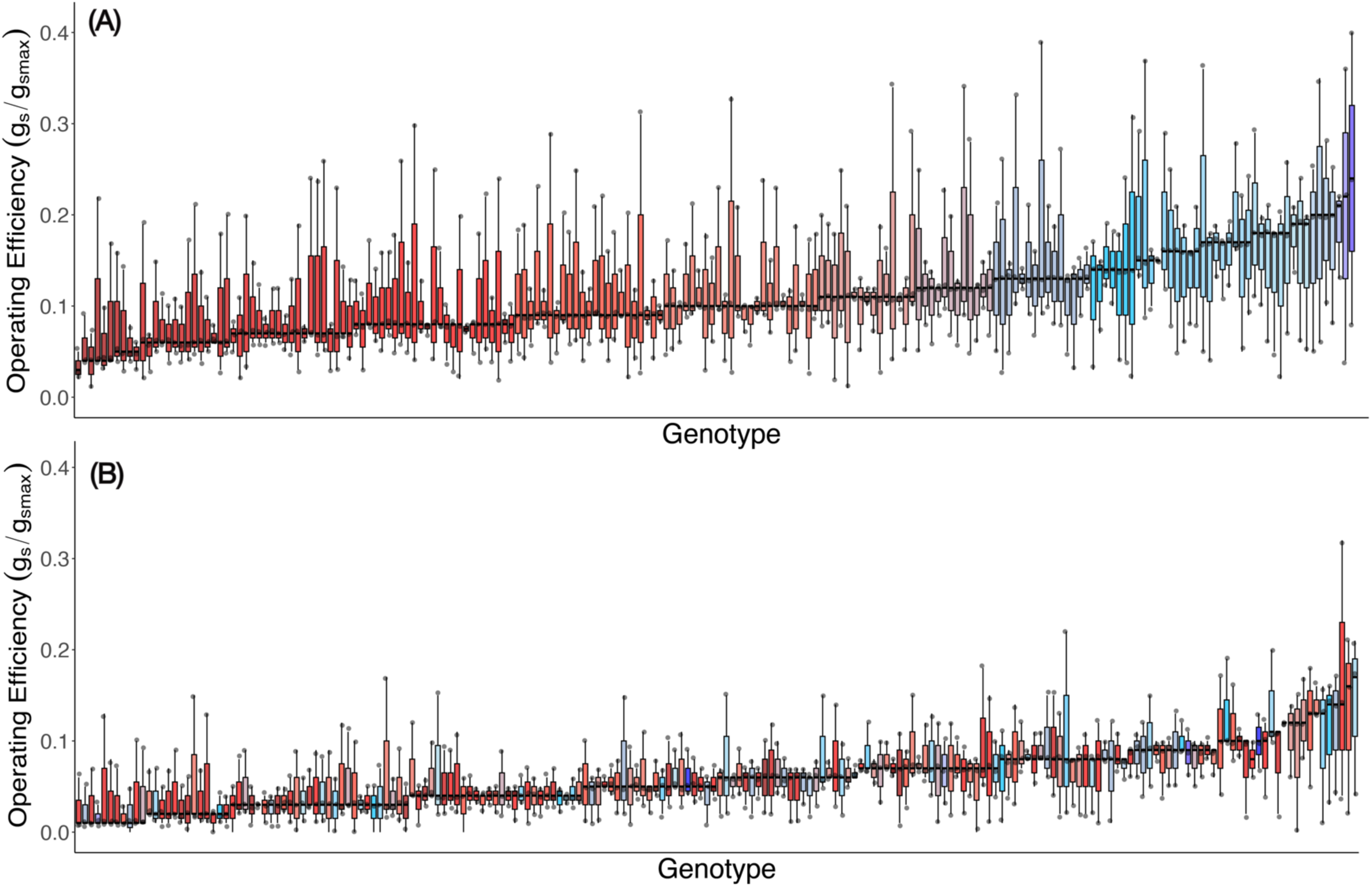
Genotypic distribution of adaxial surface stomatal conductance efficiency (*g_s_* / *g_smax_*). Data presented for 200 wheat genotypes of wheat under (A) irrigated and (B) rainfed treatments, ranked by median stomatal conductance efficiency. Colour assigned to each genotype based on irrigated stomatal conductance efficiency. Horizontal lines within boxes indicate the median and boxes indicate the upper (75%) and lower (25%) quartiles. Whiskers indicate the ranges of the minimum and maximum values. Points indicate individual measurements.

## 4 Discussion

We have successfully developed and validated a high-throughput, low-cost and open-source approach to assessing and analysing stomatal physiology and anatomy of crop species in the field. The approach comprises three steps: (1) a handheld porometer to collect *in situ* stomatal conductance measurements; (2) a handheld digital microscope, 3D printed leaf clip and integration with an open-source data collection app for non-destructive streamlined *in situ* anatomical image capture from the same leaf; and (3) a trained deep learning model to segment and analyse stomatal anatomy. The FieldDino software which was subsequently developed enables high-throughput, reliable and detailed collection of microscope images and field metadata. We have validated the use of this method on a large field trial of 200 genotypes and were able to capture the wide diversity across the wheat population we studied.

Traditional approaches to collect such data, using infra-red gas analysers (IRGAs) for stomatal physiology screening and collecting epidermal impressions for phenotyping of stomatal anatomy are not well suited to high-throughput phenotyping studies. IRGAs require up to 15 minutes for the leaf to acclimate before measurements can be taken (Taylor and Long, 2017), limiting throughput to less than 10 plants per hour (Salter *et al*., 2018). By contrast, the coupled physiology-anatomy screening approach that we have presented achieved throughput rates of over 60 plants per hour under field conditions. This has been achievable by developments in porometry and portable microscope instrumentation. There are two key reasons for the greater throughput of the porometer technique, the first of which was the speed of measurement with each sample taking < 15 s. Furthermore, the portability of the two instruments enabled fast movement from plot to plot in the field. The use of a portable handheld digital microscope gives researchers an *in situ*, non-destructive way of capturing stomatal traits at much higher throughput than conventional techniques. Each image captured using our leaf clip microscope took < 10 s per leaf. Along with increased throughput, the handheld microscopy technique is also cheaper than traditional approaches, costing less than AU $2,000 per unit.

The deep learning and computer vision scripts that we developed to analyse stomatal microscopy images were critical in enabling such a high-throughput phenotyping approach. This is supported by the conclusions of work employing similar approaches in controlled environment studies (Pathoumthong *et al*., 2023; Sun *et al*., 2023; Wang *et al*., 2024b). However, such studies have been limited to a small number of genotypes grown under glasshouse conditions as opposed to the collection of *in-situ* data from large-scale field trials. The use of deep learning for stomata anatomical analysis allows a much larger number of images and genotypes to be analysed than would be possible manually (Gibbs and Burgess, 2024). In addition to saving time, labour costs and reducing human-error, most of the deep learning model architectures for such computer vision tasks are open-access and so can easily be used and built upon in future studies. While trained models can improve image analysis throughput, model training introduces its own bottlenecks in development of training datasets requiring consistent capture and annotation. Thus, we build on the work of previous studies by developing a complete pipeline of image data collection and analysis and release the work publicly (https://github.com/williamtsalter/FieldDinoMicroscopy). Learning from our own experience with laborious image data collection, the FieldDino app bridges this high-throughput phenotyping gap, targeting these bottlenecks in the model training pipeline.

The use of porometers for the measurement of stomatal conductance has, up till now, come with a trade-off between data quality and portability/speed of measurement, certainly when compared to *g*_s_ data collected with gas exchange systems (Toro *et al*., 2019). However, the accuracy of contemporary porometers, such as the LI-COR LI-600 that we used here, is highlighted through the similarity of our data to that reported in the literature for wheat collected with gas exchange systems. The reductions in stomatal conductance that we observed under rainfed conditions of 24.4% and 28.2% for the abaxial and adaxial surfaces, respectively, were in the same order of magnitude as previously reported findings in the range of 26% - 50% under mild to moderate drought stress (Sikder *et al*., 2016; Zhao *et al*., 2020; Chen *et al*., 2022).

In addition to our stomatal conductance data being aligned with the literature, the anatomical data obtained from our approach to stomatal anatomy image collection and analysis using the deep learning model also proved to be robust and accurate. When averaged across both leaf surfaces and treatments, our model calculated the mean stomatal length to be 66.8 µm, the mean width to be 40.2 µm and the mean stomatal density to be 46.6 per mm^2^ which are of the same order of magnitude as suggested in the literature (Ricciardi *et al*., 1990; Hetherington and Woodward, 2003; Shahinnia *et al*., 2016; Wang *et al*., 2016; Faralli *et al*., 2024). The increase in stomatal density under rainfed conditions that we observed is consistent with previously reported findings (Baloch *et al*., 2013; Wang *et al*., 2016; Ouyang *et al*., 2017). Further to this, the negative correlation that we found between stomatal size and density is concurrent with previously reported findings (Dillen *et al*., 2008; Doheny-Adams *et al*., 2012; Haworth *et al*., 2023). Additionally, our finding that the adaxial leaf surface had a 31.5% higher average stomatal density than the abaxial surface and contributed more to the total stomatal conductance compared with the abaxial contributions was also consistent with previously reported results for wheat under abiotic stress conditions (Ouyang *et al*., 2017; McAusland *et al*., 2021, 2023; Wall *et al*., 2023).

Our findings support recent work which identified that a high stomatal density is advantageous under drought conditions as a way of increasing *g_smax_* to facilitate rapid gas exchange when water is readily available without causing a rise in the minimum leaf conductance, *g_leaf,min_*, which would be of detriment to drought tolerance (Ochoa *et al*., 2024). The image analysis techniques we employed here allowed us to calculate a theoretical rate of *g_smax_*, with our data aligned closely with the reported values of Pathoumthong et al., (2023), Wall et al., (2023) and Faralli et al., (2024) in the range of 0.7-2.0 mol m^-2^ s^-1^. Variations in stomatal size and density been previously reported amongst smaller populations of wheat (Faralli *et al*., 2024) and in trials investigating stomatal anatomy across a number of genotypes under drought stress (Shahinnia *et al*., 2016), but have not been captured at the scales possible using the FieldDino approach.

Combining measurements of real time stomatal physiology and anatomy as described here, allowed us to calculate the efficiency at which the stomata were operating based on their actual observed stomatal conductance and the theoretical anatomical maximum conductance, *g_smax_*. *g_sop_*relative to *g_smax_* is a key performance indicator of the tolerance of different genotypes to water limitation (Ochoa *et al*., 2024) with the operating efficiency of genotypes in our study tending to decrease under rainfed conditions compared with irrigated conditions. There is very limited research reported that has used a coupled approach to assess stomatal conductance and anatomy of the same leaf, let alone in the field (Baloch *et al*., 2013; Ouyang *et al*., 2017; Guizani *et al*., 2023). Each component of our approach has been used independently in wheat such as a porometer for conductance data collection (Sharma *et al*., 2014; Ramya *et al*., 2021; Sun *et al*., 2021, 2023), a handheld microscope for anatomical image capture (Sun *et al*., 2021, 2023; Pathoumthong *et al*., 2023) and a deep learning model for analysis of stomatal anatomy (Casado-García *et al*., 2020; Sun *et al*., 2021, 2023; Zhang *et al*., 2022; Pathoumthong *et al*., 2023). Our optimised approach incorporating all three of these techniques is a big step forwards for future research into abiotic stress tolerance, plant resource use and crop improvement.

There is an urgent need to develop crop varieties better suited to the warmer, drier, and altogether more challenging conditions our agricultural systems are likely to face in the future. This phenotyping method presents an opportunity for screening of these important traits across diverse populations in large scale, multi-environment field trials and to gain a more representative understanding of the coupled physiology-anatomy responses to stress and the genetic underpinning of these responses. With respect to yield stability, genotypes with increased stomatal conductance are better able to cope with a variety of abiotic stresses (Pooja and Munjal, 2019; Abdelhakim *et al*., 2021; Ramya *et al*., 2021). With respect to yield potential, it has already been recognised that commercial varieties have inadvertently been bred for stomatal conductance, with concurrent increases in yield occurring with yield improvement over time (Roche, 2015). It begs the question, what if we took a targeted approach where we can measure these traits at scale? Well, now we can.

### Limitations & Future Directions

Our method significantly increased throughput compared with conventional techniques while also reducing cost and improving accuracy. However, we identified several limitations to this method and discuss improvements that could be made to optimise functionality. We also address inherent limitations in the method, for example, to measure dynamic stomatal traits, that will require alternative but aligned phenotyping solutions.

In our work here, we used a 200x magnification microscope. This provided an image of a relatively large area of the leaf surface and, as such, a good sample size for measurements of stomatal density and guard cell length. However, in some of our images, the pore itself was obscured/not clear, meaning we were unable to obtain accurate direct measurements of stomatal aperture size and instead made assumptions about this based upon established, and validated, relationships between guard cell length and pore length (Franks and Farquhar, 2007; Franks and Beerling, 2009). Despite this strong relationship, this is still an assumption and we would rather measure the size of the aperture directly from collected imagery. For wheat, we would strongly advise using a 400x magnification microscope to allow direct assessment of stomatal aperture size. For other species with even smaller stomata, increased magnification may be required. For example, Pathoumthong et al. (2023) were able to accurately assess stomatal aperture with a 400x microscope for wheat but for rice and tomato, this was not possible.

Whilst we have developed some custom solutions to improve the functionality of the microscope, there were some limitations with the microscope and porometer that we were unable to overcome in this work. The leaf clip microscope mount and capture app that we developed aided focusing in the field, but the microscope was ultimately still a manual focus device. Handheld microscopes with autofocus functionality are available from Dino-Lite but at present, the resolution and magnification of these is limited. The lack of a proper light source on the LI-600 porometer also limited its functionality for measuring chlorophyll fluorescence, particularly under changeable light conditions that are often experienced in the field due to passing clouds and overstory vegetation (Durand *et al*., 2022).

For consideration in future stomatal research is the ability to measure dynamic responsiveness of stomata at scale and under field conditions. Improving the speed of stomatal responses will aid in maximising daily carbon assimilation and water use efficiency of our crop species (Lawson and Blatt, 2014). Development of robust phenotyping methods that assess stomatal responsiveness at scale are required to “complete the picture” and for inclusion of, and confidence in, stomatal traits in plant breeding programs. There is scope for the app and leaf clip we have developed and presented here to be adapted further to record videos to quantify stomatal responses in real-time.

Finally, we acknowledge that whilst the approach we present here is “high-throughput” compared to previous methodologies, it is still reliant on scientists walking around the field, plot to plot and taking measurements at the individual leaf scale. Ultimately, a truly high-throughput approach would be achieved if we were able to reliably predict these traits remotely across whole field trials, using automated ground and aerial based platforms. Thermography and hyperspectral imaging techniques present opportunities in this area (Deery *et al*., 2019; Sobejano-Paz *et al*., 2020), however, would need to be validated/ground-truthed at scales known to already work. In this sense, we could consider the stomatal phenotyping approach we present as a high-throughput validation tool for these emerging remote sensing techniques.

### Conclusions

We have developed and validated a robust, non-destructive high-throughput phenotyping approach for *in situ* coupled stomatal physiology-anatomy screening. This rapid approach allows stomatal density, stomatal size and the anatomical maximum stomatal conductance to be determined and compared to the measured operating rate of stomatal conductance, providing insights into the operating efficiency of stomata. The setup is portable and uses several ‘off-the-shelf’ products, 3D printed parts and open-source software. The versatility of this method allows it to be further adapted to assess the stomatal traits of other species using a similar equipment setup and image analysis pipelines. Compared with conventional approaches which are labour-intensive and not well suited to high-throughput *in situ* phenotyping, this screening approach takes < 15 s per measurement, making it feasible to collect informative data to assess stomatal traits across diverse populations at unprecedented scales which are relevant for plant breeding research. This is a step towards accelerating our understanding of stomatal biology within the context of yield potential and yield stability under a changing climate.

## Abbreviations

*a^’^*: Mean maximum stomatal pore area which is calculated assuming all stomatal pores are elliptical with the pore length equal to the major axis and the minor axis equal to half the pore length (υm^2^)
CAIGE: CIMMYT Australia ICARDA Germplasm Evaluation
CIMMYT: International Maize and Wheat Improvement Center
CO_2_: Carbon Dioxide
COCO: Common Objects in Context
*d*: Diffusivity of water in are at 22°C (m^2^ s^-1^)
EDPIE: Elite Diversity International Experiment
ESWYT: Elite Selection Wheat Yield Trial
F1: Harmonic mean of precision and recall
FLOPs: Floating Point Operations per Second
*g_leaf,min_*: Minimum leaf stomatal conductance
*g_s_*: Stomatal conductance
*g_smax_*: Maximum anatomical stomatal conductance
*g_sop_*: Operating rate of stomatal conductance
*g_s_*_op_/*g_smax_*: Stomatal conductance operating efficiency
H_2_O: Water
HTWYT: High Temperature Wheat Yield Trials
ICARDA: International Centre for Agricultural Research in the Dry Areas
IoU: Intersection over Union
IRGA: Infra-red gas analysers
*l*: Pore depth which is equal to the guard cell width at the centre of the stomata based on half the guard cell length (υm)
mAP: Mean average precision
Params: Parameters
ΦPSII: Quantum yield of electron transfer at PSII
PSII: Photosystem II
SAI: Stomatal Area Index
SATYN: Stress Adapted Trait Yield Nursery
SAWYT: Semi-Arid Wheat Yield Trial
SD: Stomatal density (stomata m^-2^)
*v*: Molar volume of air at 22°C (m^3^ mol^-1^)
YOLO: You Only Look Once

## Supplementary Data

The following supplementary data are available at JXB online:

**Figure S1**. Approach to stomatal annotation and regression of guard cell length to pore length relationship.

**Figure S2**. Results obtained from the testing and validation process during training of the YOLOv8 M model.

**Figure S3**. Ellipse fitting for measurement of labelled stomata.

**Figure S4**. Genotypic distribution of ΦPSII across 200 genotypes of wheat under two watering treatments.

## Acknowledgements

We thank colleagues Fiona Foster, Isobella Revell, Madalene Atkinson, Annette Tredrea and Dr Rebecca Thistlethwaite for supporting the field work in this study and Prof Richard Trethowan for supplying the wheat germplasm used for method validation.

## Author Contributions

WS: Conceptualisation; EC & WS: Data Curation; EC, GC & WS: Formal Analysis; WS: Funding Acquisition; EC: Investigation; EC & WS: Methodology; WS: Project Administration; WS: Resources; GC & WS: Software; AM & WS: Supervision; EC & WS: Validation; EC: Visualisation; EC: Writing (Original Draft); EC, GC, AM & WS: Writing – Review & Editing.

## Conflict of Interest

No conflict of interest declared

## Funding

This work was supported by funding from the Grains Research and Development Corporation (GRDC), project number ANU2304-001RTX, and the Foundation for Food and Agriculture Research (FFAR) and The International Maize and Wheat Improvement Center (CIMMYT), project number W0449.01.

## Data Availability

As outlined, all files for the 3D printed leaf clip, the Python script for stomatal annotation and the files and instructions for installing and using the FieldDino App are provided in a public GitHub repository which guides users through each step – https://github.com/williamtsalter/FieldDinoMicroscopy. Datasets used for validation of the method are available on Roboflow – https://universe.roboflow.com/narrabri-plant-physiology-hclvi/fielddino-training-set-200x.

## Tables

**Table 1:** Performance metrics used to compare different YOLOv8-seg algorithms. . Metrics used were: Parameters (Params), Floating Point Operations per Second (FLOPs), Precision (Mask), Recall (Mask), F1 Score (Harmonic mean of Precision and Recall), mAP@0.5 (mean average precision at Intersection over Union = 0.5) and mAP@[0.5:0.95] (mean average precision at Intersection over Union from 0.5 to 0.95). Based on these parameters, the YOLOv8 M model was chosen due to strong performance metrics balanced with a moderate

| Model Algorithm | Size (Pixels) | Epochs | Params | FLOPs | Precision (M) | Recall (M) | F1 Score | mAP@0.5 | mAP@[0.5:0.95] |
| --- | --- | --- | --- | --- | --- | --- | --- | --- | --- |
| YOLOv8n | 640 | 315 | 3.4 | 12.6 | 0.9630 | 0.9292 | 0.9458 | 0.9659 | 0.6401 |
| YOLOv8s | 640 | 115 | 11.8 | 42.6 | 0.9633 | 0.9239 | 0.9432 | 0.9659 | 0.6424 |
| YOLOv8m | 640 | 326 | 27.3 | 110.2 | 0.9651 | 0.9377 | 0.9512 | 0.9709 | 0.6592 |
| YOLOv8l | 640 | 456 | 46 | 220.5 | 0.9651 | 0.9360 | 0.9503 | 0.9700 | 0.6762 |
| YOLOv8x | 640 | 239 | 71.8 | 344.1 | 0.9665 | 0.9376 | 0.9518 | 0.9718 | 0.6682 |

model size.

## Notes

### Competing Interest Statement

The authors have declared no competing interest.

